# Multi-Domain Translation between Single-Cell Imaging and Sequencing Data using Autoencoders

**DOI:** 10.1101/2019.12.13.875922

**Authors:** Karren Dai Yang, Anastasiya Belyaeva, Saradha Venkatachalapathy, Karthik Damodaran, Adityanarayanan Radhakrishnan, Abigail Katcoff, GV Shivashankar, Caroline Uhler

## Abstract

The development of single-cell methods for capturing different data modalities including imaging and sequencing have revolutionized our ability to identify heterogeneous cell states. While various methods have been proposed to integrate different sequencing data modalities, coupling imaging and sequencing has been an open challenge. We here present an approach for integrating vastly different modalities by learning a probabilistic coupling between the different data modalities using autoencoders to map to a shared latent space. We validate this approach by integrating single-cell RNA-seq and chromatin images to identify distinct sub-populations of human naive CD4+ T-cells that are poised for activation. Collectively, our approach provides a framework to integrate and translate between data modalities that cannot yet be measured within the same cell for diverse applications in biomedical discovery.

---

Recent evidence has highlighted the importance of the 3D organization of the genome to regulate cell-type specific gene expression programs (*1, 2*). High-throughput and high-content single-cell technologies have provided important insights into genome architecture (using imaging and chromosome capture methods) (*3–5*) as well as detailed genome-wide epigenetic profiles and expression maps (using various sequencing methods) (*6–8*). However, high-throughput paired measurements of these different data modalities within single cells is still a major challenge requiring significant breakthroughs in single-cell technologies. We here present a computational framework for integrating and translating between different data modalities such as imaging and sequencing which cannot yet be obtained experimentally in the same cell, thereby providing a methodology to predict the genome-wide expression profile of a particular cell given its chromatin organization and vice-versa. Such a methodology is valuable to understand how features in one dataset translate to features in the other.

Different data modalities provide different perspectives on a population of cells and their integration is critical for studying cellular heterogeneity and its function (Fig 1a). Current computational methods allow the integration of datasets of the same modality (*9–11*) or of different modalities with the same data structure such as various sequencing measurements (*12, 13*). To integrate and translate between data modalities with very distinct structures, we propose a new strategy of mapping each dataset to a shared latent representation of the cells (Fig 1b). This mapping is achieved using autoencoders (*14–16*), neural networks consisting of an encoder (mapping to the latent space) and a decoder (mapping back to the original space), whose architectures can be customized to the specific data modality (Fig 1b-c). Combining the encoder and decoder modules of different autoencoders enables translating between different data modalities at the single-cell level (Fig 1d). To enforce proper alignment of the embeddings obtained by the different autoencoders, we employ a *discriminative objective function* to ensure that the data distributions are integrated in the latent space, and, when prior knowledge is available, additional objective functions that encourage the alignment between specific markers or the anchoring of certain cells. For an in-depth discussion of the alignment strategy, see Materials and Methods and (*17*). While our method is designed to integrate vastly different data structures, in Fig S1 (see also Table S1, S2) we show that our framework is competitive with previous methods (based on canonical correlation analysis (*18*)) for the simpler problem of integrating different modalities with similar data structures, namely RNA-seq and ATAC-seq from (*19*).

**Figure 1:**
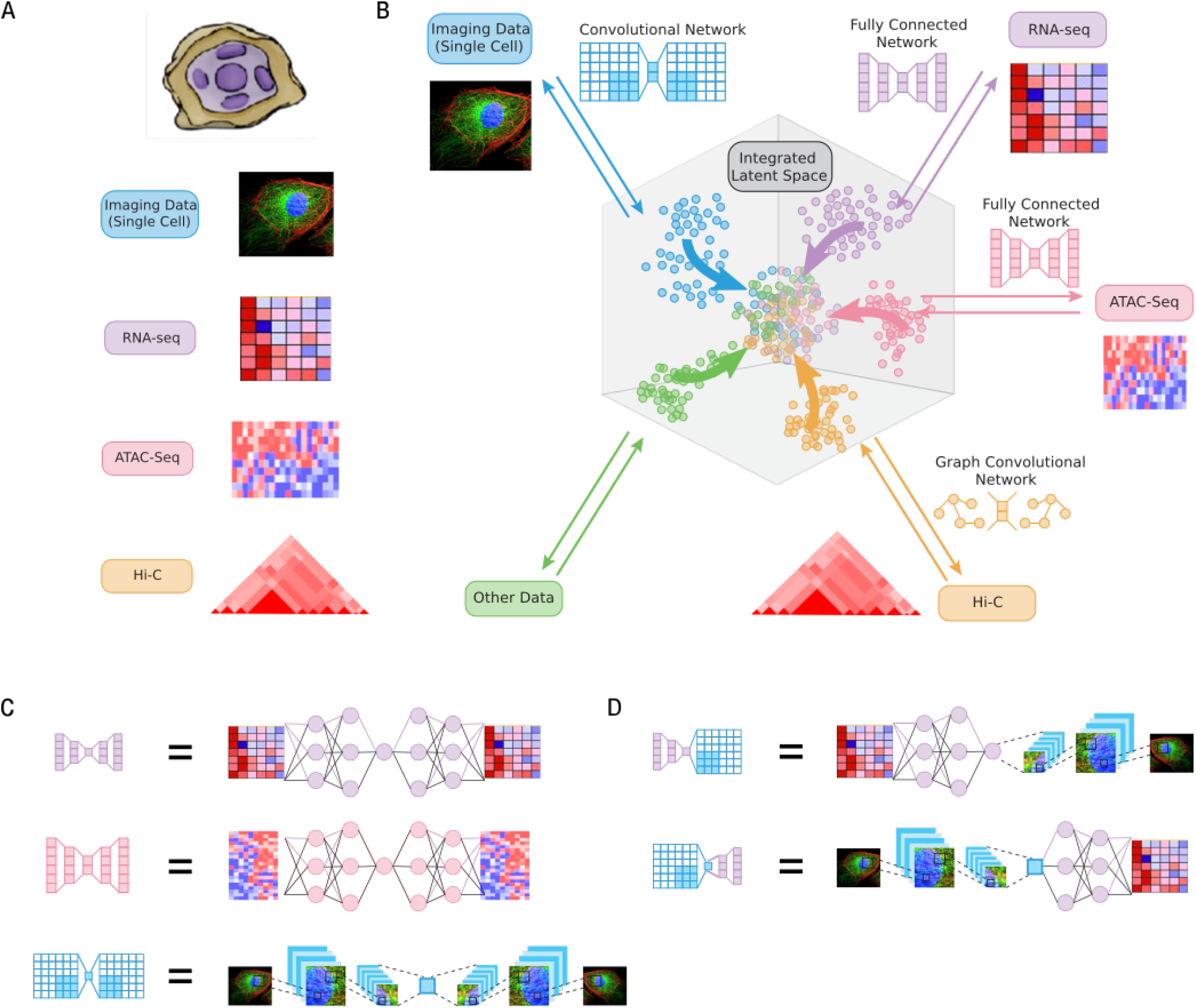
Schematic of multimodal data integration and translation strategy using autoen-coders. (a) Each modality or dataset presents a different view of the same underlying population of cells of interest. (b) Our computational strategy to integrate multiple modalities involves embedding each dataset into a shared space that represents the latent state of the cells, such that the distributions of each dataset mapped into the latent space are aligned. (c) The embedding of each dataset is performed using an autoencoder, a neural network with separate encoder and decoder modules, whose architectures can be customized to the specific data modality. (d) Combining the encoder and decoder modules of different autoencoders enables translation between different data modalities at the single-cell level.

We applied our method to integrate single-cell RNA-seq data with chromatin images in order to study the heterogeneity within naive T-cells. T-cell activation is a fundamental biological process and identifying naive T-cells poised for activation is critical to understanding immune response (*20*). Moreover, linking genome organization with gene expression generates hypotheses that can be tested experimentally to validate our methodology. We first analyzed single-cell RNA-seq data of human blood cells from (*21*) and used known markers to identify naive (CD4+) and activated T-cells (Fig 2a and Fig S2, Table S3, Data S1). An in-depth analysis of the naive T-cell population revealed two distinct subpopulations that were robust to various clustering strategies (Fig 2b and Fig S3, Data S1). Differential gene expression and GO enrichment analysis indicated that one cluster corresponded to quiescent cells while the other was poised for activation, with an expression profile more similar to that of activated T-cells (Fig 2c-d).

**Figure 2:**
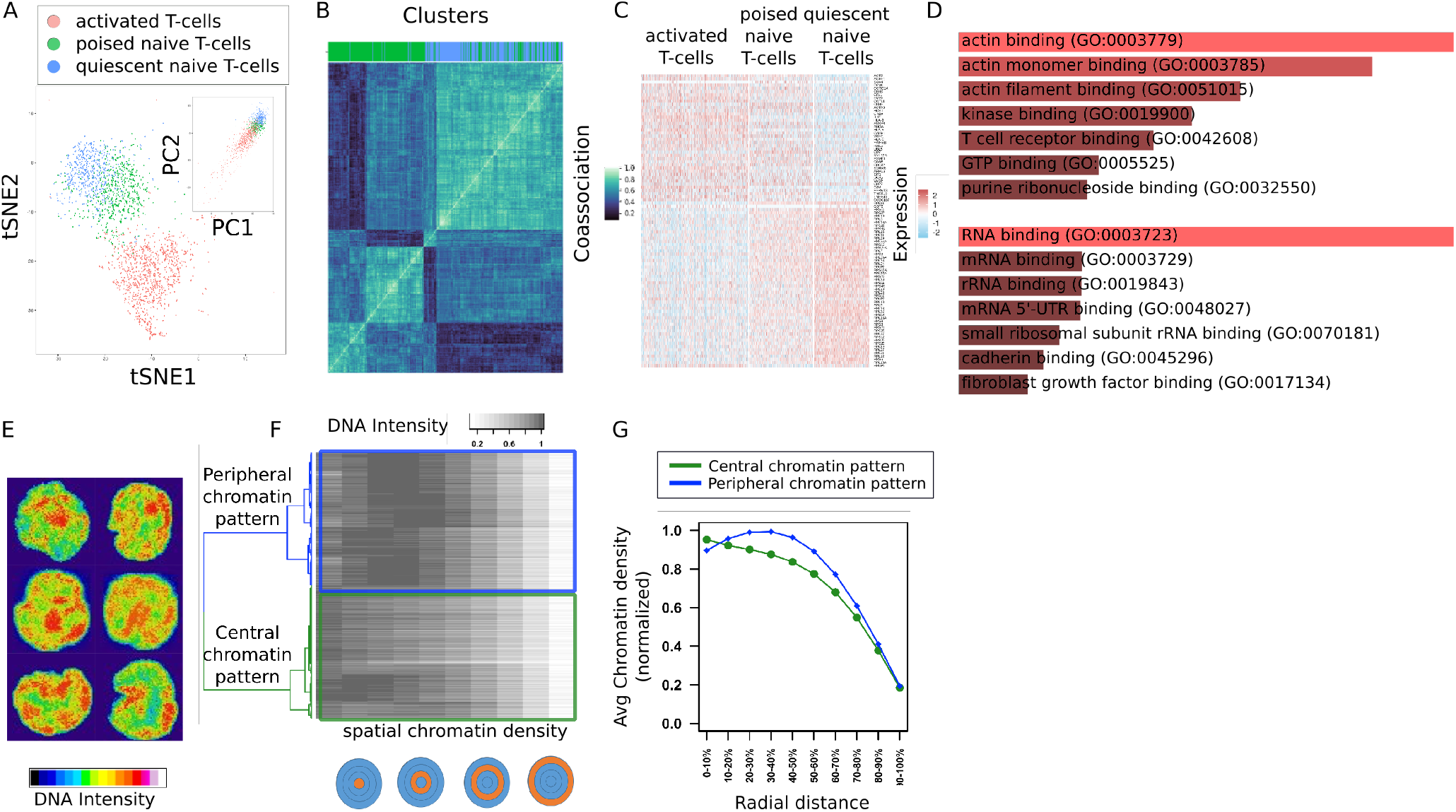
Analysis of single-cell RNA-seq data and single-cell chromatin images of naive CD4+ T-cells reveals two distinct subpopulations respectively. (a) tSNE and PCA (inset) embeddings of single-cell RNA-seq data from (*21*). Cluster analysis reveals activated (red) population of T-cells and naive population of T-cells divided into two subpopulations (blue and green). (b) Consensus clustering plot demonstrating the robustness of quiescent and poised clusters of naive T-cells to various clustering methods (c) Differential gene expression analysis between the blue and green subpopulations reveals two distinct gene expression programs. The green subpopulation of naive T-cells is more similar to the activated T-cells and hence poised for activation, while the blue subpopulation shows an upregulation of ribosomal genes and has a relatively more quiescent expression profile. (d) GO enrichment analysis of marker genes for quiescent and poised naive T-cell subpopulations supports two distinct gene expression programs. (e) Examples of DAPI-stained nuclear images of naive CD4+ T-cells. (f) Cluster analysis of the 3D nuclear images is performed by first quantifying the chromatin signal in concentric spheres with increasing radii, and then using hierarchical clustering on these spatial chromatin features. (g) One cluster has higher concentration of chromatin in the central region of the nucleus (green), while the other cluster has higher concentration of chromatin in the peripheral region of the nucleus (blue).

Given the link between expression and chromatin organization (*22*), we hypothesised the presence of two subpopulations of naive T-cells with distinct chromatin packing features. To test this, we carried out DAPI-stained imaging experiments of naive CD4+ human T-cells and analyzed their chromatin organization (Fig 2e and Fig S4). We extracted image features by quantifying the chromatin density in concentric spheres with increasing radii (Fig 2f). Cluster analysis based on the extracted features revealed two distinct subpopulations of cells, with higher chromatin density in the central and peripheral nuclear regions respectively (Fig 2g). These observations are consistent with previous experiments in mouse naive T-cells that also showed two subpopulations with distinct chromatin organization patterns, where naive T-cells with more central heterochromatin were shown to be poised for activation (*23*).

Up to this point, we had observed two subpopulations of naive T-cells based on a separate analysis of gene expression (from single-cell RNA-seq data) and chromatin packing (from single-cell imaging data). To link the identified subpopluations from the unpaired datasets, we used our method to integrate the single-cell RNA-seq data with the chromatin images (see Materials and Methods and Table S4), thereby enabling translation between the two data modalities (Fig 3a and Fig S5). To assess whether our methodology aligns imaging features and gene expression features in a consistent manner, we analyzed the latent embeddings as well as the results of translation between the two datasets. Visualization of the latent representations revealed that the subpopulations from the two datasets were appropriately matched (Fig 3b and Fig S6). In addition, we found that classifiers trained to distinguish between the subpopulations in the original datasets also performed well when evaluated on the translated datasets (Fig 3c). Importantly, in the gene expression matrix that was translated from the imaging dataset, we found that the differential expression of genes was strongly correlated with the true observed differential expression of genes in the real RNA-seq dataset and that the predicted and observed enriched gene sets were highly overlapping (Fig 3d-e).

**Figure 3:**
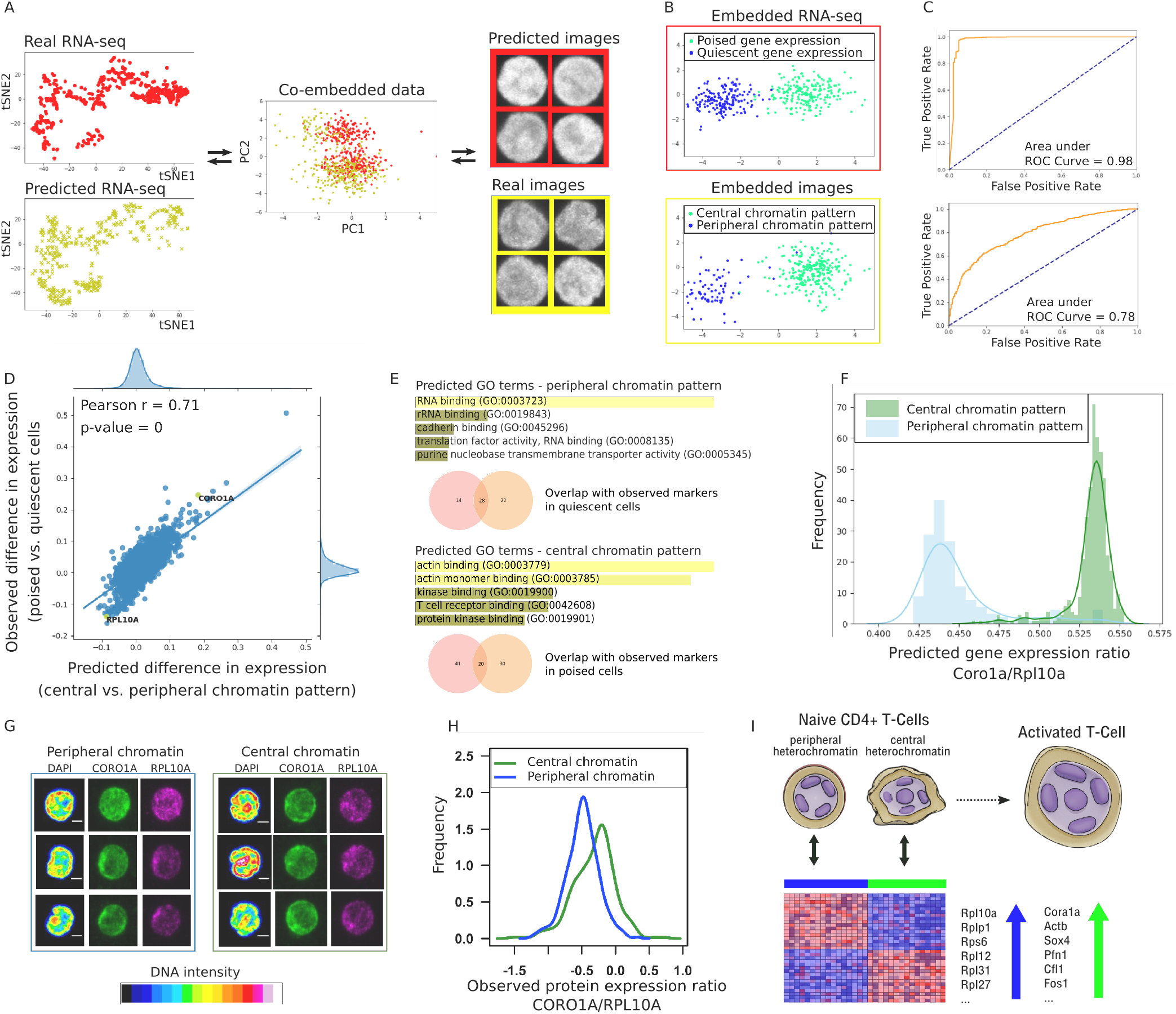
Integration of single-cell RNA-seq data and single-cell nuclear images of naive T-cells using our methodology allows translating between chromatin packing and gene expression profiles. (a) Illustration of data integration and translation: (left) tSNE plots of observed single-cell RNA-seq data (red) and single-cell RNA-seq data translated from single-cell images (yellow); (middle) PCA visualization of single-cell RNA-seq data (red) and single-cell imaging data (yellow) embedded in 128-dimensional latent space; (right) examples of observed single-cell images (yellow) and images translated from single-cell RNA-seq data (red). (b-e) Evidence that our data integration methodology correctly aligns gene expression features and imaging features. (b) Linear Discriminant Analysis (LDA) plots of single-cell RNA-seq (top) and imaging (bottom) datasets embedded in the latent space. The clusters with more quiescent (blue) and poised (green) gene expression programs from the RNA-seq dataset are aligned with the clusters with peripheral (blue) and central (green) chromatin patterns from the imaging dataset. (c) (Top) Receiver Operating Characteristic (ROC) curve illustrating performance of a classifier trained to distinguish between peripheral and central chromatin patterns in images when evaluated on images translated from RNA-seq data. (Bottom) ROC curve illustrating performance of a classifier trained to distinguish between quiescent and poised gene expression programs when evaluated on RNA-seq data translated from images. High performance of both classifiers indicates that the alignment of the clusters in the latent space in (b) also holds in the original gene expression and imaging spaces. (d) Differential gene expression analysis between cells with central and peripheral chromatin pattern performed on the predicted gene expression matrix translated from images using our methodology. The predicted fold-change of gene expression based on images is strongly correlated with the observed fold-change of gene expression between quiescent and poised naive T-cells from the actual RNA-seq dataset. (e) Analysis of GO enrichment terms of cells with central and peripheral chromatin pattern based on the predicted gene expression matrix translated from images using our methodology shows a high overlap between predicted markers from the imaging dataset and actual markers from the RNA-seq dataset. (f-h) Validation of our model alignment using single-cell immunofluorescence experiments. (f) Histograms of predicted Coro1a/Rpl10a gene expression ratio in cells with central (green) and peripheral (blue) chromatin pattern based on the gene expression matrix translated from the imaging dataset. Our model predicts the upregulation of Coro1a and Rpl10a in the cells with central and peripheral chromatin pattern respectively. (g) Examples of immunofluorescence staining data of CORO1A and RPL10A proteins collected along with the chromatin images. (h) Histograms of measured CORO1A/RPL10A protein ratio in cells with central (green) and peripheral (blue) chromatin pattern. Consistent with the model prediction, CORO1A and RPL10A proteins are upregulated in the cells with central and peripheral chro-matin pattern respectively. (i) Schematic of the two naive T-cell subpopulations characterized by our multimodal analysis, in which peripheral and central patterns of chromatin density are associated with gene expression programs for quiescent and poised naive CD4+ T-cells respectively.

Our model generates predictions of gene expression programs based on patterns of chromatin density (Fig 3e). To validate these results experimentally, we chose two genes, Coro1a and Rpl10a, which are predicted to be strongly upregulated in the naive T-cell subpopulations with peripheral and central patterns of chromatin density respectively (Fig 3d,f). We analyzed the immunofluorescence staining data of these proteins obtained along with chromatin images (Fig 3g). Consistent with the model predictions, we found that CORO1A was upregulated in the cells with central chromatin pattern, while RPL10A was upregulated in the images with peripheral chromatin pattern (Fig 3h and Fig S7). These results altogether demonstrate that our method properly aligns the gene expression and image features that characterize two distinct sub-populations of human naive T-cells, and suggest that peripheral and central enrichment of chromatin are associated with gene expression programs for more quiescent and poised naive CD4+ T-cells respectively (Fig 3i).

In summary, we presented a powerful approach to integrate and translate between different data modalities of very different structures, namely single-cell chromatin imaging and RNA-seq. Using our methodology we quantitatively analyzed the link between chromatin organization and expression, thereby identifying a subpopulation of naive T-cells, which is poised for activation. Importantly, our methodology can be applied generally to integrate single-cell datasets that cannot yet be measured in the same cell, and as such has broad implications for the integration of spatial transcriptomics (*24*), proteomics (*25*) and metabolomics (*26*) datasets. In particular, this methodology can be applied to predict the functional landscape of single cells in a tissue slice where only limited functional data is available by acquiring chromatin imaging data.

## Acknowledgments

The authors thank Diego Pitta de Araujo for the schematic drawings. K.D.Y. was supported by the National Science Foundation (NSF) Graduate Research Fellowship and ONR (N00014-18-1-2765). A.B. was supported by J-WAFS and J-Clinic for Machine Learning and Health at MIT. The G.V.S. laboratory thanks the Mechanobiology Institute (MBI), National University of Singapore (NUS), Singapore and the Ministry of Education (MOE) Tier-3 Grant Program for funding. A.R. was supported by the National Science Foundation (DMS-1651995). C.U. was partially supported by NSF (DMS-1651995), ONR (N00014-17-1-2147 and N00014-18-1-2765), a Sloan Fellowship, and a Simons Investigator Award. The Titan Xp used for this research was donated by the NVIDIA Corporation.

## Data and code availability

The primary images and the code will be made available upon acceptance of the manuscript.

