## Supplementary Materials for "Multi-Domain Translation between Single-Cell Imaging and Sequencing Data using Autoencoders"

**This PDF file includes:**

Materials and Methods
Figures S1 to S7
Tables S1 to S4
Captions for Data S1
References

**Other Supplementary Materials for this manuscript include the following:**

Data S1 [Cluster labels for PBMCs, Cluster labels for naive CD4+ T-cells, Differential expression poised vs. quiescent]

### Materials and Methods

#### Multi-domain data integration and translation with autoencoders

In multi-modal data integration, each modality or dataset presents a different view of the same underlying population of cells. We formalize this problem within a probabilistic framework. Formally, we consider cells from each dataset as samples of random variables $X_1, X_2, \ldots, X_k$ that are generated independently from a common latent variable $Z$:

$$X_i = f_i(Z, N_i), \quad \forall i = 1 \cdots k,$$

where $f_i$ are functions, $Z$ is a latent variable with distribution $P_Z$, and $N_i$ are noise variables. The domain of $Z$ represents some underlying latent representation space of cell state, and each function $f_i$ represents a random function through which data sampled from $X_i$ is generated. For simplicity of notation, we assume in the remainder of the discussion that $X_i$ is a deterministic function of $Z$, so that the noise variables can be ignored. This model implies the following factorization of the joint distribution of data $P_{\mathbf{X}}$:

$$p_{\mathbf{X}}(\mathbf{x}) = \int_{\mathcal{Z}} \Pi_{i=1}^{k} p_{X_i|Z}(x_i|z) p_Z(z) dz, \tag{1}$$

where $p_Z$ is the probability density of $Z$, and $p_{X_i|Z}$ is the conditional distribution of $X_i$ given $Z$ that reflects the generative process. Multi-modal data integration can then formalized as the problem of learning conditional distributions $p_{X_i|Z}$ as well as latent distribution $p_Z$ based on samples from the marginal distributions $P_{X_1}, P_{X_2}, \ldots P_{X_k}$, which are given by our datasets.

When the latent distribution $P_Z$ is known, then learning the conditional distributions $p_{X_i|Z}$ between the marginals $P_{X_1}, P_{X_2}, \ldots, P_{X_k}$ can be solved by learning multiple autoencoders. Specifically, for each domain $i = 1, \ldots, k$, we propose training a regularized encoder-decoder pair $(E_i, D_i)$ to minimize:

$$\mathbb{E}_{x \sim P_{X_i}} \left[ \lambda_1 L_1(x, D_i(E_i(x))) + \lambda_2 L_2(E_i \# P_{X_i} | P_Z) \right], \tag{2}$$

where $\lambda > 0$ is a hyperparameter, $L_1$ is the (Euclidean) distance metric, $L_2$ represents a divergence between probability distributions, and $E_i \# P_{X_i}$ is the distribution of $X_i$ after embedding to the latent space $\mathcal{Z}$. Translation from domain $i$ to $j$ is accomplished by composing the encoder from the source domain with the decoder from the target domain, i.e. taking

$$X_{i \to j}(x_i) := D_j(E_i(x_i)). \tag{3}$$

The autoencoders obtained by minimizing the above loss satisfy various consistency properties (see [17]). Since $P_Z$ is not usually known in practice, it must also be estimated from the data using one of two approaches: (1) learn $P_Z$ by training a regularized autoencoder on data from a single representative domain; or (2) alternate between training multiple autoencoders until they agree on an invariant latent distribution. The first approach is typically more stable to train in practice, while the second captures variability across multiple domains and is therefore more suitable for integrating multiple datasets. Note that $P_Z$ is by no means unique; there are multiple solutions that can result in the same observed data distributions.

Concretely, we can learn an invariant latent distribution based on two domains $i, j \in \{1, \ldots, k\}$ as follows. Let $\hat{P}_{Z_{i'}}, i' \in \{i, j\}$ denote the empirical latent distribution based on encoded data from domain $i'$, i.e. $\hat{P}_{Z_{i'}} = E_{i'} \# P_{\mathcal{X}_{i'}}$. Then for domain $i$, we optimize the objective,

$$\min_{E_i, D_i} \mathbb{E}_{x \sim P_{X_i}} L_1(x, D_i \circ E_i(x)) + \lambda L_2(E_i \# P_{X_i} | P_{\hat{Z}_j}),$$

while for domain $j$, we optimize the objective,

$$\min_{E_j, D_j} \mathbb{E}_{x \sim P_{X_j}} L_1(x, D_j \circ E_j(x)) + \lambda L_2(E_j \# P_{X_j} | P_{\hat{Z}_i}).$$

In practice, we parameterize $(E_i, D_i)$ using neural networks and minimize the objective function in (2) via stochastic gradient updates. In particular, we can choose $L_2$ to be the *discriminative* loss,

$$L_2(P|Q) := \max_f \mathbb{E}_{x \sim P} \log f(x) + \mathbb{E}_{x \sim Q} \log(1 - f(x)),$$

which is equivalent to the Jensen-Shannon divergence up to a constant factor.

Prior knowledge is sometimes available to guide the integration of different data modalities. For example, there may be knowledge of alignment of specific markers or clusters, or knowledge of certain samples from different datasets corresponding to the same cell, i.e. the same point in the latent space. In this case, training of the autoencoders can be guided by additional loss functions that incorporate the prior knowledge.

**Shared markers/clusters from both datasets.** If there are shared markers or clusters that are present in two datasets, they can be aligned by replacing $L_2$ above with the following discriminative loss that is conditioned on these factors:

$$L_2(P|Q) := \max_f \mathbb{E}_{x,y\sim P} \log f(x,y) + \mathbb{E}_{x,y\sim Q} \log(1 - f(x,y)), \quad (4)$$

where $P$ and $Q$ are now joint distributions over the data and the markers and/or clusters. Alternatively, if the markers or clusters have $k$ discrete values (i.e. $1, \ldots, k$), then we can add a simple classifier model $p_\theta(Y|Z)$ with parameters $\theta$ and minimize the loss

$$\sum_{\text{modality } i} \mathbb{E}_{x,y\sim P_i} \sum_{j=1}^{k} \mathbb{1}(y = j) p_\theta(Y = j | Z = E_i(x)) \quad (5)$$

with respect to $\theta$ and the parameters of the encoders $E_i$; here $P_i$ is the distribution of the $i$th data modality.

**Anchored cells in both datasets.** If $(x_1, x'_1), (x_2, x'_2), \cdots, (x_m, x'_m)$ are anchored points from two datasets that are embedded by encoders $E, E'$, we can add the following *anchor* loss,

$$\sum_{i=1}^{m} ||E(x_i) - E'(x'_i)|| \quad (6)$$

to minimize their distance in the latent embedding space.

#### Model validation on paired RNA-seq and ATAC-seq data

We obtained paired RNA-seq and ATAC-seq data collected in the same cell from [19]. Specifically, we used paired data collected from human lung adenocarcinoma–derived A549 cells treated with dexamethasone (DEX) for $0, 1$, or 3 hours. We downloaded the single-cell RNA-seq data from the GEO accession number GSE117089, corresponding to [19]. For the ATAC-seq data, instead of using raw matrix of peaks by cells, we acquired a transcription factor (TF) motif by cells matrix from the authors, which was computed as described in [19] by counting occurrences of each motif in all accessible sites for each cell, resulting in 815 TF motifs. For single-cell RNA-seq data we considered genes that were determined to be differentially expressed by [19], keeping genes with q-value $> 0.05$. Both single-cell RNA-seq and ATAC-seq were $\log(x + 1)$ transformed and normalized to zero mean and unit variance. The number of cells that were shared between TFs $\times$ cells matrix from ATAC-seq and genes $\times$ cells matrix from RNA-seq was 1874, therefore our model based on paired autoencoders had to learn a latent embedding and translate between data sets with different number of features, i.e. for single-cell RNA-seq a matrix of 2613 genes $\times$ 1874 cells and for single-cell ATAC-seq a matrix of 815 TFs $\times$ 1874 cells.

We trained paired autoencoders to embed single-cell RNA-seq and ATAC-seq into the same latent space (dimensionality of 50), which allows mapping and translation of samples from one space to the other. Our model's architecture consisted of fully connected layers with input, hidden layers and output sizes listed in Table S1. In order to train the model we minimized the weighted sum of losses listed in Table S2. The model was trained in Pytorch with learning rate of 0.0001 and batch size of 32 for 4000 epochs using Adam with $\beta_1 = 0.5, \beta_2 = 0.999$ and weight decay of 0.0001.

Since the RNA-seq and ATAC-seq data was collected in the same cell, we could evaluate the accuracy of our method in matching samples from RNA-seq to ATAC-seq (and vice-versa). For evaluation, we created an 80-20 training-test split of the paired data. We used two different metrics, calculated on the test set, to measure the accuracy of mapping RNA-seq and ATAC-seq samples in the latent space. For both metrics, we used $\ell_1$ distance for distance computations in the latent space.

- First we considered the $k$-nearest neighbors accuracy between the projections of each data set into the latent space

$$\text{k-NN(A, B)} = \frac{\sum_i \mathbb{1}(b'_i \in a^k_i)}{n}, \quad (7)$$

where $n$ is the length of the test set, $A$ and $B$ are sets of vectors, with $b_i$ as a vector in $B$ and $a_i$ as its pair in A, and $b_i'$ and $a_i'$ are the encoded versions of $b_i$ and $a_i$ in the latent space. The set $a_i^k$ contains the $k$ nearest neighbors of $a_i'$ in $A'$, the set of vectors in $A$ projected into the latent space. Since k-NN(A, B) does not necessarily equal k-NN(B, A), we computed the average of these metrics.

- We also computed the fraction of samples closer than the true match: More precisely, for each encoded point from domain $A$, we know its true nearest neighbor in domain $B$ since the data is paired. After encoding both domains (RNA-seq and ATAC-seq) into a shared latent space, we computed the distance (in the inferred latent space) between a point in domain $A$ and its true match in domain $B$. We then computed the fraction of points in the test set that are closer in distance than the true match. The fraction is averaged over all data points and translation directions (A to B and B to A). For perfectly paired data, this fraction would be 0.

We compared our method against deep canonical correlation analysis (DCCA), which uses paired samples between two domains to learn a shared embedding of the two domains by maximizing the total correlation [18]. The model for DCCA consisted of two neural networks, one for each domain. For ATAC-seq data, the input to the model was a matrix with 815 features, followed by 815 hidden nodes with sigmoid activation, and a final output layer of size 50. For RNA-seq data, the input to the model was a matrix with 2613 features, followed by 2613 hidden nodes with sigmoid activation, and a final output layer of size 50. Finally, as in [18], linear CCA was applied to the output layers of the two neural networks corresponding to the two different domains. DCCA jointly learns the parameters for both neural networks such that the correlation of the final output layer between the domains is maximized. DCCA was trained using RMSProp with learning rate of $10^{-3}$, batch size of 1024 for 100 epochs. Regularization parameter of $10^{-9}$ was applied to the networks.

For both the autoencoder (our method) and DCCA, we explored including supervision, i.e. points that are paired between the two domains (anchored cells in both data sets). For supervision, we included an additional term in the loss function corresponding to mean absolute error between paired training points in the latent space. Our method based on autoencoders does not require paired samples, however DCCA does. In order to train DCCA with 0% supervision, we used treatment time labels of the cells to generate paired data. For each point with a particular treatment time label, we sampled 100 random points with the same label to use as its paired samples.

Fig. S1 shows that our method using autoencoders performs at least as well as DCCA for integrating single-cell RNA-seq and single-cell ATAC-seq data.

#### Gene expression data of naive CD4+ T-cells

The gene expression data corresponsding to human peripheral blood mononuclear cells (PBMCs) was collected by [21] and filtered cell by gene matrix was downloaded from `https://support.10xgenomics.com/single-cell-gene-expression/datasets/2.1.0/pbmc8k`. We analyzed the PBMC 8k data set since it had the highest number of reads per cell. Since the data was already filtered we applied minor additional filtering such as removing cells with high proportion of counts in mitochondrial genes ($\geq 10\%$), which reduced the number of cells from 8381 to 8371 cells. After cell filtering, we performed gene filtering by removing mitochondrial genes and keeping genes which had at least 10 cells expressing the gene with a count $> 1$, resulting in 7633 remaining genes.

After cell and gene filtering, we followed a standard analysis pipeline using Seurat (version 2.3.0) [9, 12]. We normalized the gene expression measurements for each cell by the total expression for that cell and scaled the result using the median total expression across cells. The scaled result was $\log(x + 1)$ transformed. We z-scored the data and applied PCA to obtain 30 components, which were used for t-SNE and clustering analysis. The t-SNE embedding for all cells, computed using default parameters, is shown in Fig. S2a. We clustered the data using default clustering in Seurat (FindClusters function) with resolution parameter of 0.4, which resulted in 13 clusters, shown in Fig. S2a. Briefly, the clustering method in Seurat constructs a $k$-nearest neighbor graph and adjusts the edge weights between cells based on Jaccard similarity of their local neighborhoods. The resulting graph is clustered using the Louvain algorithm to obtain cell clusters. In order to determine the identity of each cluster we performed differential expression analysis using the default Wilcoxon rank sum test in Seurat (FindAllMarkers function). We list the top 10 differentially expressed genes for each cluster in Table S3.

From the clustering analysis of all PBMCs and annotation using differentially expressed genes, we were able to determine that cluster 1 generally corresponds to naive CD4+ T-cells (differential overexpression of CCR7, LEF1, TCF7), cluster 2 corresponds to cytotoxic T-cells (differential overexpression of GZMK, NKG7, CCL5), cluster 3 corresponds to activated CD4+ T-cells (differentially overexpression of IL7R, IL32) and cluster 4 corresponds to naive CD8+ T-cells (differentially overexpression of CD8A, CD8B, LEF1, CCR7) [27, 28]. Fig. S2b provides a t-SNE plot of all PBMCs, colored by the expression of known markers genes, further corroborating our cell type annotation.

#### Gene expression analysis of naive CD4+ T-cells

We aimed to explore potential heterogeneity in naive CD4+ T-cell gene expression in relation to CD4+ T-cells that already underwent activation. We performed a feature selection step, keeping genes which had average log-fold change of $> 0.05$ between naive and activated CD4+ T-cells (and vice versa), resulting in 1187 genes. Similar to the analysis of PBMCs, we applied PCA for dimensionality reduction on the selected genes, keeping the top 30 components and clustered the naive CD4+ T-cells using the default clustering in Seurat with resolution of 0.8. Fig. S2c shows the resulting clusters. Based on differential expression analysis and t-SNE embedding, the smallest cluster (shown in grey in Fig. S2c) was determined to belong to the CD8+ T-cell population since the top differentially overexpressed genes for this small cluster were CD8A and CD8B. Therefore, this small cluster was removed from the further gene expression analysis of the naive CD4+ T-cells. In order to characterize the remaining two subpopulations, we performed differential expression analysis on the two subpopulations of naive CD4+ T-cells using Wilcoxon rank sum test. We defined marker genes as all genes with Bonferroni-corrected p-value of $< 0.05$. Fig 2c, shows the resulting heatmap for the genes that are markers between poised and quiescent subpopulations of naive T-cells and are also part of the 1187 genes that have an average log-fold change of $> 0.05$ between naive and activated CD4+ T-cells (and vice versa). Gene ontology analysis was performed on these marker genes overexpressed in each cluster (average log fold change$> 0$) using Enrichr [29, 30], keeping gene ontology molecular function terms with lowest p-values.

#### Cluster robustness

Our clustering is robust across different number of clusters and clustering methods. We re-clustered the data corresponding to naive CD4+ T-cells using Seurat with different resolution parameters, i.e. $0.9, 1.1$ and $1.15$ to obtain $3, 4$ and $5$ clusters respectively. We computed the silhouette coefficient for each clustering (Fig. S3a), observing that the number of clusters corresponding to 2 gave the highest score. This suggests that using 2 clusters is optimal. We also fit a Gaussian mixture model to the data and computed the Bayesian information criterion (BIC) for a model with $1, 2, 3, 4$ and $5$ mixture components (across 100 randomly initialized trials). As shown in Fig. S3b, the model with 2 components resulted in the lowest mean BIC, suggesting again that 2 clusters is optimal for this data.

In order to check robustness to different clustering methodologies, we also used k-means, Gaussian mixture models and spectral clustering based on $k$-nearest neighbor graph with $k \in \{10, 20, 50, 100\}$ to cluster the data. We performed 100 different initializations for each method and computed the co-association matrix, which quantifies how often each pair of cells was clustered together using a particular method; the result is shown in Fig. 2b. We observe that the chosen clustering given by Seurat is in agreement with the co-association matrix.

#### Cell Culture and immunostaning

CD4+/CD45RA+ naive helper T-cells from human peripheral blood were purchased from AllCells. These cells were revived and cultured in media (RPMI-1640 + 10% FBS + 1% pen-strep) as per the manufacturer's instructions. The cells for the experiments were used within two days upon revival.

Cells in media were allowed to adhere to Poly-lysine coated slides for 30 minutes. Cells were then fixed with 4% Paraformaldehyde (Sigma) for 30 minutes and washed with PBS three times, which also removed unattached cells. Permeabilization was done with 0.5% Triton X-100 (Sigma) for 10 minutes followed by PBS washes. Blocking was done with 5% BSA in PBS for 30 minutes and incubated with primary and secondary antibodies as per the dilution and incubation time recommended by the manufacturer. Cells were washed with PBS (+0.1% Tween) three times after primary and secondary antibody incubation. During the final step, excess liquid was removed by slanting the slides. ProLong® Gold Antifade Mountant with DAPI (ThermoFischer Scientific) was added to these slides and allowed to cure for 24 hours. Coverslips were then sealed and imaged using a confocal microscope.

#### Confocal Microscopy and Image Analysis

1024 × 1024 and 12-bit multi-channel images were obtained using a Nikon A1R confocal microscope. Z-stack images were captured using a 100× objective with a pixel size of 0.1 $\mu$m and 0.5 $\mu$m depth. Images were processed and further analyzed using custom programs in Fiji and R.

The nuclear boundaries were segmented in 3D using the DAPI channels to identify individual nuclei. These nuclei were eroded by 0.5 microns in $x, y$, and $z$ iteratively until the volume of the eroded nucleus was less than 10 cubic microns. Then the mean intensity of each 3D ring (width 0.5 microns) in the nucleus was computed for all cells. The intensity fraction was calculated by normalizing the mean ring intensity for each nucleus (maximum= 1). Linear interpolation was then used to compute the intensity fraction of rings that occupy 0-10% to 90-100% volume fraction of the nucleus. The heatmaps were visualized using functions from gplots, RColorBrewer and dendextend.

In order to calculate the cellular levels of proteins, the 3D nuclear object was dilated by 2 microns in $x, y$ and $z$. This was efficient as the cells were all spherically shaped with high karyoplasmic index. The total intensity in the 3D cellular object was computed for each protein channel and their ratio was obtained for each cell.

#### Model training on single-cell RNA-seq data and single-cell chromatin images

Since the imaging dataset is more difficult to reconstruct in comparison to the RNA-seq dataset, we first pretrained the image autoencoder to reconstruct single-cell chromatin images for 850 epochs using the reconstruction loss and the cluster classifier loss (Equation 5). Subsequently, we trained the full model consisting of the pretrained image autoencoder, the RNA-seq autoencoder, and latent space discriminator using reconstruction loss and conditional discriminative loss with hyperparameters $\lambda_1 = 0.1, \lambda_2 = 1$. The architectures of all networks are shown in Table S4. Models were trained with the Adam optimizer with a learning rate of 1e-3. Images were normalized to range between $[0, 1]$ and RNA-seq matrix was $\log(x+1)$ normalized.

#### Receiver Operating Characteristic (ROC) analysis on translated data sets

In order to assess whether translated image and RNA-seq data sets are able to still separate poised and quiescent subpopulations as well as central and peripheral subpopulations based on features from the original data sets and analyze if the clusters obtained separately from gene expression and imaging data sets align with each other, we performed ROC analysis on the translated data sets. For RNA-seq, we trained a random forest classifier (using 100 trees in a forest with 2 as the maximum depth of a tree) on reconstructed RNA-seq data with labels based on poised versus quiescent clustering of naive CD4+ T-cell gene expression data. This classifier essentially learned the genes that separate the two clusters. Next, we translated chromatin images into RNA-seq using our autoencoder method and assessed the performance of the pre-trained classifier on its ability to separate central versus peripheral clusters on images translated to RNA-seq (ROC analysis in Fig. 3c, top).

Similarly, to assess translation of RNA-seq into images, we trained a classifier (using architecture similar to the image encoder in Table S4) to separate central versus peripheral chromatin patterns. Then, we translated RNA-seq data into images and evaluated the performance of the pre-trained classifier in being able to separate poised versus quiescent clusters (ROC analysis in Fig. 3c, bottom). The area under the curve (AUC) was computed for both of these tasks, resulting in high AUCs for both translation directions.

#### Differential expression analysis on images translated to RNA-seq

Imaging data sets can provide a rich quantification of cells, such as their chromatin organization. Based on image analysis, subpopulations of cells with different characteristics may be found (e.g. central versus peripheral chromatin organization), and it is often of interest to study which genes might be markers of each subpopulation such that these subpopulations can be separated for example using antibodies against the marker genes. However, generally the full gene expression and imaging features cannot be measured in the same cell. Our computational framework can translate chromatin images into RNA-seq and calculate the predicted mean difference in expression between the subpopulations (e.g. central versus peripheral chromatin organization). As shown in Fig. 3d, the observed mean difference in expression is strongly correlated with the predicted mean difference in expression.

For images translated to RNA-seq, we obtained a set of marker genes associated with central and peripheral chromatin organization by performing Welch's t-test on the generated RNA-seq data (considering marker genes for each cluster to be the top 50 genes that had the highest mean difference in expression for the two clusters as well as p-value $< 0.05$ after adjustment for multiple hypothesis testing using the Benjamini–Hochberg procedure). We performed gene ontology analysis on the marker genes for each cluster and in Fig. 3e we report the 5 gene ontology molecular function terms with lowest p-values.

#### Autoencoder training on chromatin images

We trained a convolutional autoencoder with the following architecture on the chromatin images: (1) We used 15 convolutional layers with 256 $3\times3$ filters per layer followed by leaky ReLU activations throughout; (2) Layers 2-6 have a stride size of 2 and Layers 8-12 are followed by bilinear upsampling layers with a scale factor of 2. The bottleneck of our network thus provides a 256 dimensional representation of the images. We trained our network using the Adam optimizer (learning rate of $10^{-4}$) and used a Kaiming uniform initialization for all our convolutional layers. All of the images were trimmed to remove background and resized to $32 \times 32$ images in order to remove nucleus size as a distinguishing feature. We held out 10% of the data as test data and trained until the reconstruction loss on the test data was smaller than $10^{-3}$.

In order to determine whether our network was able to separate the poised and quiescent naive CD4+ T-cell clusters (as determined by the protein ratio of CORO1A to RPL10A) in an unsupervised fashion, we visualized the embedding of the images corresponding to the histogram peaks in Fig. 3h (namely the images with protein ratio in the range [.64, .7] and [0.93, 1]). Fig. S7 shows the resulting t-SNE embedding, where the color coding corresponds to the protein ratio of CORO1A to RPL10A. Interestingly, the latent embedding of the images obtained in an unsupervised fashion (with no information about the proteins) captures the protein ratio.

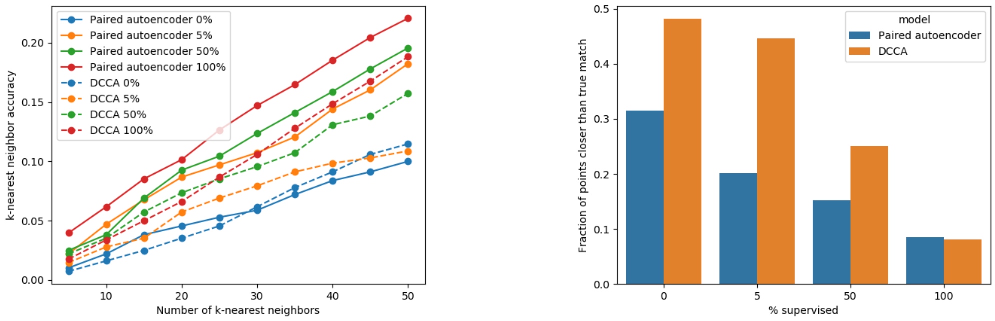


(a) (b)

Fig. S1: Performance of our autoenocoder method and DCCA on paired RNA-seq and ATAC-seq data. (a) $k$-nearest neighbor accuracy for quantifying the quality of matching between local neighborhoods. Autoencoder and DCCA were trained with $0, 5, 50$ and 100% supervision (samples that are known to be paired). (b) Fraction of points closer than the true match for DCCA and autoencoder models with $0, 5, 50$ and 100% supervision.

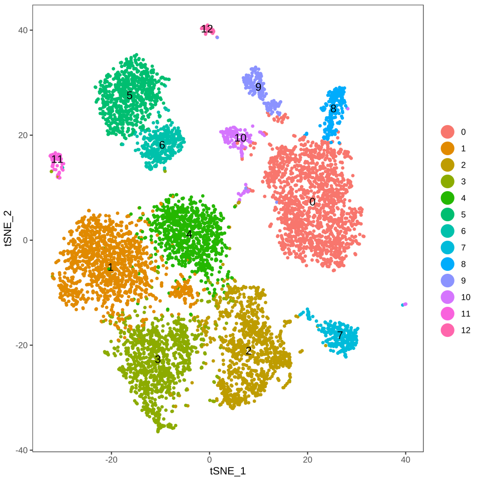


(a)

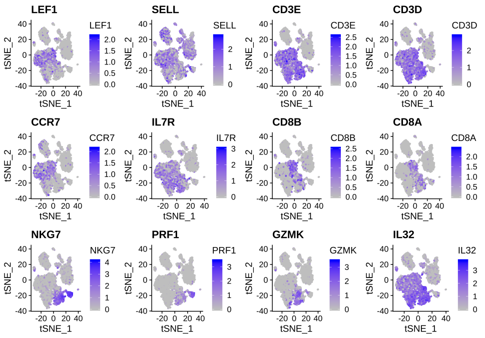


(b)

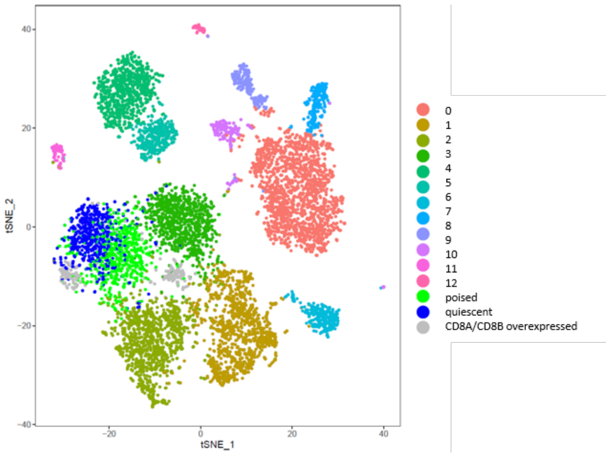


(c)

Fig. S2: Clustering of PBMC data. (a) t-SNE of all cells in the PBMC data set, colored by inferred cluster label. (b) t-SNE plots of all cells in the PBMC data set, colored by expression of genes marking naive T-cell subpopulations (LEF1, SELL, CCR7), T-cells (CD3E, CD3D, IL7R, IL32), CD8 T-cells (CD8A, CD8B), natural killer, and cytotoxic T-cells (NKG7, PRF1, GZMK). (c) t-SNE plot including the clustering of the naive CD4+ T-cells. Grey subpopulation differentially overexpresses CD8A and CD8B as the genes with highest average log-fold change (corrected p-value = $1.30 \times 10^{-40}$ and $1.54 \times 10^{-51}$ respectively), indicating that these cells are not naive CD4+ T-cells; thus they have been removed from further analysis.

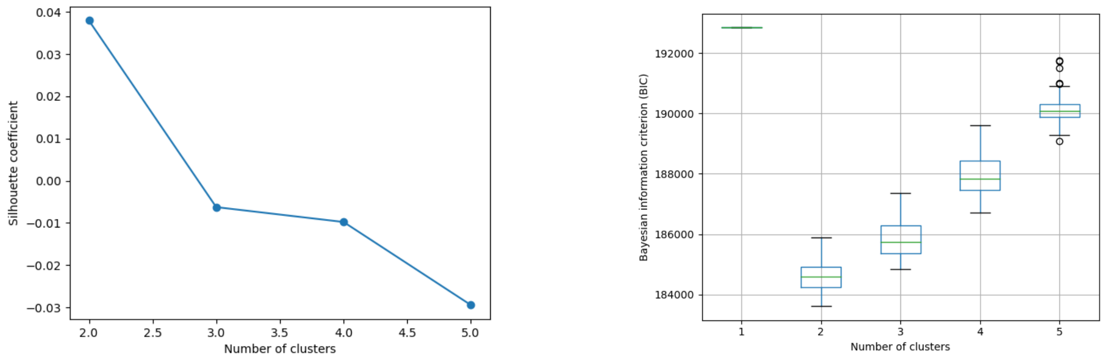


(a) (b)

Fig. S3: Evaluating optimal number of clusters for naive CD4+ T-cell gene expression data. (a) Silhouette coefficient for clusters obtained with Seurat at different resolutions (0.8, 0,9, 1.1, 1.15). (b) BIC score (averaged over 100 trials) for Gaussian mixture model with 1, 2, 3, 4 and 5 components.

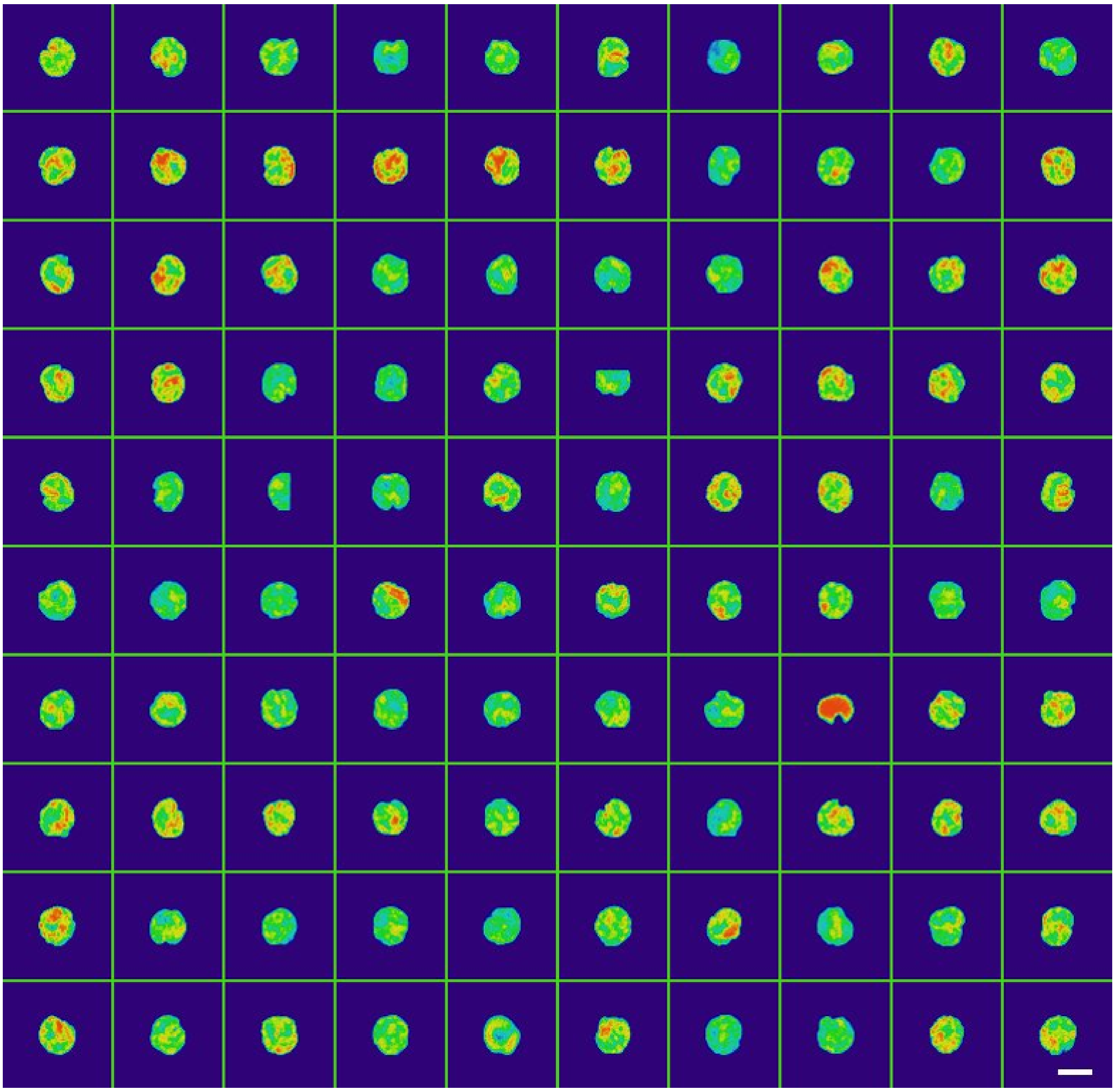

Fig. S4: Examples of naive CD4+ T-cell nuclei stained with DAPI. Scale bar is 2 microns.

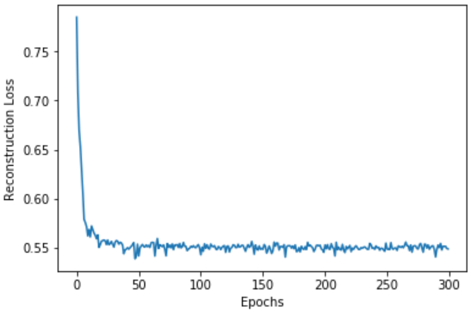


(a)

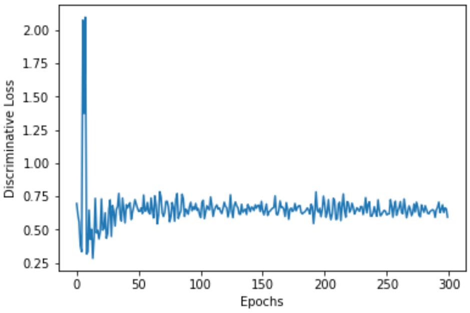


(b)

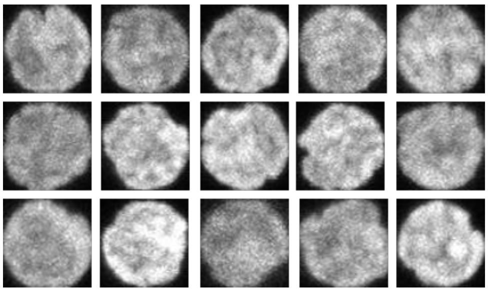

(c)

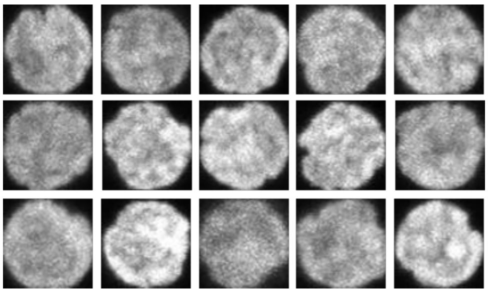

(d)

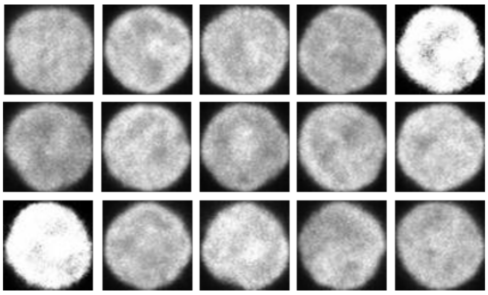

(e)

Fig. S5: Model trained on single-cell RNA-seq and single-cell images of DAPI-stained nuclei. (a) Reconstruction loss curve (sum of RNA-seq and image reconstruction losses). (b) Discriminative loss curve for RNA-seq and image translation model. (c) Examples of input images to the image autoencoder. (d) Reconstructed images after training the image autoencoder. (e) Generated images translated from RNA-seq to image space.

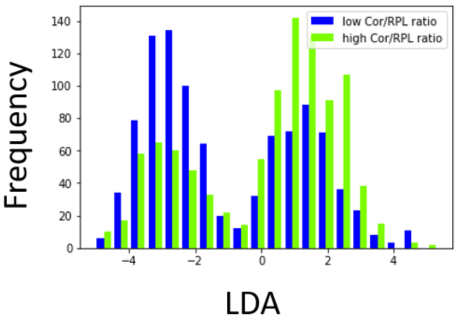


(a)

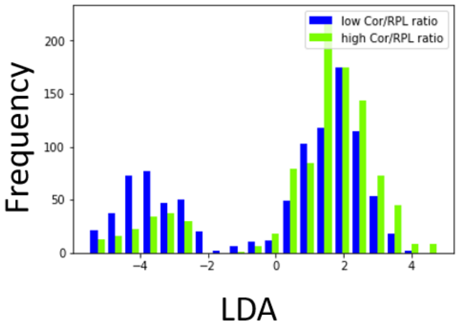


(b)

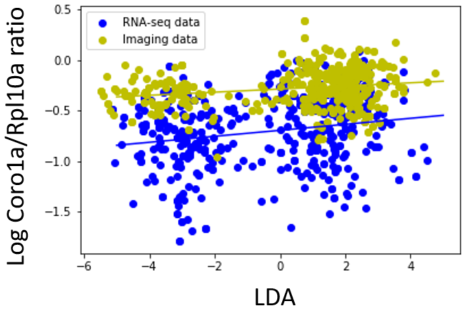


(c)

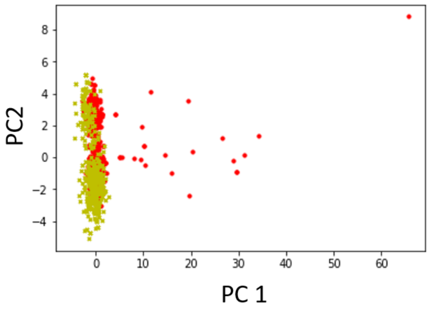


(d)

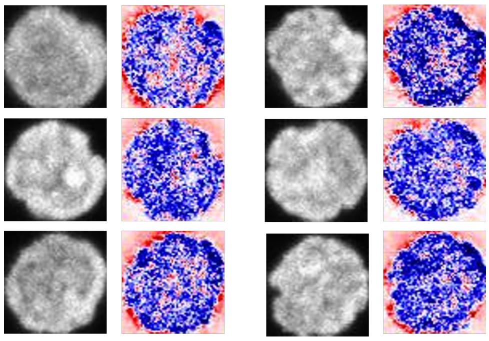

(e)

Fig. S6: Validation of inferred latent embedding. Histograms of embedded naive CD4+ T-cells from (a) RNA-seq and (b) imaging data sets, split by high versus low CORO1A/RPL10A ratio. Histogram is computed along LDA axis that maximally separates two subpopulations in the latent space, showing that the axis aligns with CORO1A/RPL10A ratio. (c) Scatterplot of CORO1A/RPL10A ratio versus projection onto LDA axis. In both data sets, the positive correlation between the ratio and the projection onto the LDA axis is statistically significant (p-value $< 10^{-5}$). (d) RNA-seq (red) and imaging (yellow) data embedded in latent space, visualized using PCA. (e) Interpretation of image features along the LDA axis that maximally separates the two naive T-cell subpopulations in the latent space. Results show decreased background chromatin concentration in the nucleus.

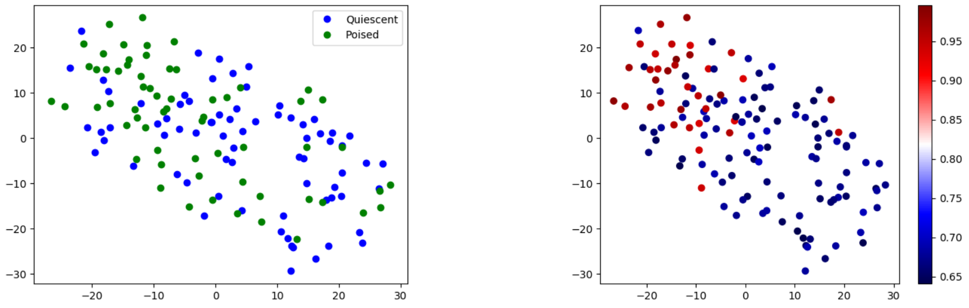


(a) (b)

Fig. S7: t-SNE visualization of latent space for convolutional autoencoder colored by (a) cluster label and (b) protein ratio of CORO1A to RPL10A. The autoencoder separates out the two naive CD4+ T-cell clusters by protein ratio without cluster supervision.

| | Input size | Hidden layer size(s) | Output size |
|---|---|---|---|
| **Encoder A** | 815 (ATAC-seq TFs) | 815, 815, 815, 100 | 50 (latent space) |
| **Encoder B** | 2613 (RNA-seq genes) | 2613, 2613, 2613, 100 | 50 (latent space) |
| **Decoder A** | 50 (latent space) | 100, 815, 815, 815 | 815 (ATAC-seq TFs) |
| **Decoder B** | 50 (latent space) | 100, 2613, 2613, 2613 | 2613 (RNA-seq genes) |
| **Discriminator** | 50 (latent space) | 50, 100 | 1 |
| **Classifier** | 50 (latent space) | N/A | 3 (treatment time class probabilities) |

Table S1: Network architecture for autoencoder network trained on RNA-seq and ATAC-seq data collected from A549 cells. The discriminator, decoders, and encoders have Leaky ReLU activations after each layer.

| Loss description | Type | Weight |
|---|---|---|
| Reconstruction loss for ATAC-seq | Mean absolute error | 10 |
| Reconstruction loss for RNA-seq | Mean absolute error | 10 |
| Discriminative loss | Mean squared error | 10 |
| Shared cluster (treatment time) classification loss for ATAC-seq | Cross-entropy | 10 |
| Shared cluster (treatment time) classification loss for RNA-seq | Cross-entropy | 10 |
| Anchor/supervision loss between paired points in the latent space | Mean absolute error | 0.1 |

Table S2: Losses and corresponding weights for autoencoder network trained on RNA-seq and ATAC-seq data collected from A549 cells.

| Cluster # | Differentially overexpressed genes | Cluster annotation |
|---|---|---|
| 0 | S100A8, S100A9, LYZ, S100A12, TYROBP, FCN1, FTL, CTSS, MNDA, CST3 | |
| 1 | LDHB, CCR7, LEF1, RPL31, NOSIP, CD3E, RPS27, RPS6, SARAF, TCF7 | Naive CD4+ T-cells |
| 2 | CCL5, NKG7, GZMK, GZMA, IL32, KLRB1, CST7, DUSP2, CMC1, CTSW | Cytotoxic T-cells |
| 3 | IL32, LTB, IL7R, ITGB1, KLRB1, LDHB, CD3D, CD2, AQP3, GSTK1 | Activated CD4+ T-cells |
| 4 | CD8B, CD8A, JUNB, LDHB, LEF1, CCR7, NPM1, RPS6, CD7, SARAF | Naive CD8+ T-cells |
| 5 | TCL1A, CD79A, CD74, CD79B, MS4A1, HLA-DRA, HLA-DPA1, HLA-DQB1, HLA-DPB1, CD37 | |
| 6 | CD79A, MS4A1, CD79B, CD74, JCHAIN, HLA-DRA, HLA-DPA1, HLA-DPB1, HLA-DQB1, BANK1 | |
| 7 | GNLY, NKG7, PRF1, FGFBP2, GZMA, CTSW, KLRD1, GZMB, KLRF1, SPON2 | |
| 8 | LST1, FCGR3A, AIF1, SAT1, FCER1G, COTL1, PSAP, MS4A7, FTL, IFITM3 | |
| 9 | HLA-DQA1, HLA-DRB1, CST3, HLA-DPB1, HLA-DPA1, FCER1A, HLA-DRA, CD74, HLA-DQB1, LYZ | |
| 10 | PPBP, PF4, GNG11, HIST1H2AC, RGS18, TUBB1, TSC22D1, S100A9, S100A8, NRGN | |
| 11 | CD79A, CD79B, CD74, TCL1A, MS4A1, CD37, BANK1, HLA-DPA1, CD22, RALGPS2 | |
| 12 | GZMB, JCHAIN, LILRA4, ITM2C, PTGDS, IRF7, IRF8, PLD4, PLAC8, CCDC50 | |

Table S3: Top 10 differentially upregulated genes (average log-fold change $> 0$) for each cluster in PBMC data set.)

| | Image autoencoder | RNA-seq autoencoder |
|---|---|---|
| Encoder | 2D Convolutional Block (1, 128, 4 x 4, 2) | Fully connected block (7633, 1024) |
| | 2D Convolutional Block (128, 256, 4 x 4, 2) | Fully connected block (1024, 1024) |
| | 2D Convolutional Block (256, 512, 4 x 4, 2) | Fully connected block (1024, 1024) |
| | 2D Convolutional Block (512, 1024, 4 x 4, 2) | Fully connected block (1024, 1024) |
| | 2D Convolutional Block (1024, 1024, 4 x 4, 2) | Fully connected block (1024, 1024) |
| | Fully connected (4096, 128) | Fully connected (1024, 128) |
| Decoder | Fully connected (128, 4096) | Fully connected (128, 1024) |
| | 2D Transposed Convolutional Block (1024, 1024, 4 x 4, 2) | Fully connected block (1024, 1024) |
| | 2D Transposed Convolutional Block (1024, 512, 4 x 4, 2) | Fully connected block (1024, 1024) |
| | 2D Transposed Convolutional Block (512, 256, 4 x 4, 2) | Fully connected block (1024, 1024) |
| | 2D Transposed Convolutional Block (256, 128, 4 x 4, 2) | Fully connected block (1024, 1024) |
| | 2D Transposed Convolutional Block (128, 1, 4 x 4, 2) | Fully connected (1024, 7633) |
| | Sigmoid | |

Table S4: Network architecture for RNA-seq and image autoencoder networks. Each block consists of a batch normalization layer and ReLU nonlinearity. The discriminator has the same structure as the RNA-seq decoder with no batch normalization, 3 fully connected blocks and output dimension of 2.

Data S1: Cluster label assignments based on single-cell RNA-seq for PBMC cells. Cluster label assignment for naive CD4+ T-cells based on single-cell RNA-seq. Differential expression of genes between quiescent and poised naive CD4+ T-cells.
